# Preliminary evidence for altered interoceptive processing in smokers

**DOI:** 10.1101/529842

**Authors:** Jennifer Todd, Farah Hina, Jane E Aspell

**Author notes:** Corresponding Author: Miss Jennifer Todd.

## Abstract

**Background:** Neuroimaging evidence suggests that interoceptive processing might be altered in nicotine addiction, however this has not yet been confirmed with behavioral measures. Therefore, we investigated the perception of internal bodily states in smokers (49) and people who had never smoked (n=51), by measuring interoceptive sensitivity (IS) and interoceptive awareness (IA).

**Methods:** IS was measured with a heartbeat tracking task and a heartbeat discrimination task. Performance on the heartbeat tracking task may be influenced by one's ability to estimate an elapsed time interval so this was controlled by a time-estimation (TE) task. IA was measured using two sub-scales from the Multidimensional Assessment of Interoceptive Awareness (MAIA). Participants in the ‘addiction’ group completed the Revised Fagerström Test for Nicotine Dependence (FTND-R) to measure addiction severity.

**Results:** Independent-samples *t*-tests revealed that non-smokers performed significantly better than smokers on the heartbeat tracking task (*p* = 0.007, *d* = 0.22). There were no group differences observed for the remaining variables. Furthermore, none of the variables predicted addiction severity.

**Conclusions:** This is the first demonstration of behavioral differences in interoception in participants with nicotine addiction. When considered in the context of previous research, these findings support the hypothesis that interoceptive processing may be disturbed in addiction. These data also support the hypothesis that behavioral and self-report measures of interoception assess two separate constructs.

---

Cigarette smoking modifies several bodily states, for example it causes persistent increases in blood pressure and heart rate (Groppelli, Giorgi, Omboni, Parati & Mancia, 1992), and has powerful sensory effects on the airways (Westman, Behm & Rose, 1996). Furthermore, nicotine withdrawal symptoms include decreased heart rate and gastrointestinal discomfort (Kenny & Markou, 2001). The processing of such internal bodily sensations is termed *‘*interoception*’*. *Interoception* refers to a collection of processes by which the physiological condition of the body is communicated to the brain: afferent signals are received, processed, and integrated to directly or indirectly affect the behavior of an organism, with or without conscious awareness (Cameron, 2002; Craig, 2002). There is a growing body of evidence to suggest that interoception is a multi-dimensional construct (Garfinkel, Seth, Barrett, Suzuki & Critchley, 2015). Interoceptive *sensitivity* (IS) describes the objective detection of internal bodily sensations with behavioral measures such as heartbeat perception tasks (e.g. Schandry, 1981; Whitehead, Drescher, Heiman & Blackwell, 1977). Meanwhile, interoceptive *awareness* (IA; also referred to as ‘interoceptive sensibility’ —see Garfinkel et al. 2015) refers to self-reported (via questionnaires) awareness of internal bodily sensations (e.g. Mehling et al 2012; Porges, 1993). Neuroimaging studies have identified the insular cortex as a crucial region for interoceptive processing (Craig, 2002; Critchley, Weins, Rotshtein, Öhman & Dolan, 2004).

The several reasons for hypothesizing that disturbed interoceptive processing is a crucial component of smoking addiction. Firstly, research suggests other forms of addiction are associated with disturbed interoception. Substance addictions have been related to altered structure and functioning of the insula (Paulus & Stewart, 2014). Moreover, neuroimaging studies have related substance addictions to the exaggerated processing of both aversive (Berk *et al.,* 2015) and pleasant (Stewart, May, Tapert & Paulus, 2015) non-drug related interoceptive stimuli. Stewart *et al.* (2015) additionally observed that the processing of interoceptive feeling states may change over the course of an addiction: participants who had recently developed a stimulant addiction showed exaggerated neural processing of non-drug related pleasant interoceptive feeling states compared to participants with a chronic drug addiction.

Secondly, a relationship between the insula and smoking has been identified repeatedly, although there is some debate as to how this relationship impacts the IS of smokers. Some studies suggest that drug addiction sensitizes the insula, which would predict enhanced IS in smokers. For example, Naqvi, Rudrauf, Damasio and Bechara (2007) observed that smokers with brain damage involving the insula were significantly more likely than smokers with brain damage not involving the insula to undergo a ‘disruption of smoking addiction’ (p. 532) – that is, the ability to cease smoking with ease, immediately, and without persistent cravings or relapses. The insula has subsequently been associated with cue-induced nicotine cravings (Engelmann *et al*., 2012). In particular, the strength of connectivity between the right anterior insula and the precuneus (which has been associated with interoception and self-awareness), is positively correlated with the magnitude of craving responses to smoking cues (*r^2^* = .15, *p* <.01; Moran-Santa Maria *et al*., 2015). Furthermore, within the human cortex, the density of nicotinic acetylcholine receptors is highest in the insular and anterior cingulate cortices (Picard *et al.,* 2013). In contrast, more recent neuroimaging studies have been interpreted to suggest that nicotine addiction is associated with reduced interoceptive signaling within the insular cortex, which would therefore predict diminished IS in smokers (Droutman, Read & Bechara, 2015). For example, smoking has been associated with significant reductions in the thickness of the right insula (Morales, Ghahremani, Kohno, Hellemann & London, 2014), and nicotine dependence scores have been shown to negatively correlate with gray-matter volume of the bilateral anterior insula (Wang *et al.,* 2019). Furthermore, smoking has been associated with reduced resting state functional connectivity between the right anterior insula and several other areas including the anterior cingulate cortex, ventromedial prefrontal cortex, amygdala, left dorsolateral prefrontal cortex and dorsal striatum (Bi *et al.,* 2016). Meanwhile, other evidence suggests that IS may fluctuate with levels of satiation/withdrawal (Avery *et al.,* 2016).

Despite evidence from neuroimaging studies supporting the involvement of the insula (and by implication, interoceptive processing) in nicotine addiction, there is a paucity of research investigating the roles of IS and IA in addiction, and no studies have examined whether there are differences between nicotine-addicted participants and healthy controls. Ateş Çöl, Sonmez and Vardar (2016) found that participants with an addiction to alcohol displayed lower levels of IS (measured by the ‘heartbeat tracking task’; Schandry, 1981) in comparison to healthy, control participants. However, there was a significant difference between the smoking habits of the two experimental groups, with a greater proportion of individuals in the ‘alcohol-addicted’ group also addicted to smoking cigarettes. It is therefore conceivable that the differences between the two conditions were caused by a combination of alcohol and smoking addictions, or exclusively by an addiction to smoking. Meanwhile, Sönmez *et al.* (2016) found that participants with an addiction to either alcohol, heroin, or synthetic cannabinoids also demonstrated lower levels of IS (measured with the heartbeat tracking task), in comparison to healthy controls. However, significant differences in smoking habits were also present, and additionally, participants were undergoing inpatient treatment for addiction (including pharmacological therapies that may affect bodily states) at the point of testing. In contrast, a recent study conducted by Betka et al. (2018; smoking status of participants unknown) found that heavy (non-clinical) alcohol consumption was not associated with IS. Together, the findings from these studies suggest that addiction specifically, rather than non-clinical consumption of a substance, is associated with impaired IS. However, further research is required to confirm this hypothesis, and whether this is applicable to nicotine addiction.

Research considering interoceptive ability has focused mainly on cardiac detection, and there is an implicit assumption that cardiac detection (possibly due to the frequent adoption of such tasks) represents a marker of ‘general’ interoceptive ability (Tsakiris & Critchley, 2016). Specifically, the heartbeat tracking task (Schandry, 1981) and the heartbeat discrimination task (Brener & Kluvitse, 1988; Katkin, Reed & Deroo, 1983; Whitehead *et al.*, 1977) are most commonly utilised (Tsakiris & Critchley, 2016). The tracking task requires participants to count the number of heartbeats experienced over discrete time intervals, and these numbers are compared to the actual number of cardiac events. Meanwhile, the discrimination task requires participants to listen to a series of tones, and identify whether they are synchronous or asynchronous with their own cardiac rhythm. These tasks are generally considered to be both valid and reliable, with good test-retest reliability (Knoll & Hodapp, 1992). However, psychometric weaknesses have also been acknowledged (Tsakiris & Critchley, 2016). The heartbeat tracking task is susceptible to participants’ abilities to estimate time, and their knowledge/expectations about their own heart rate (Knapp-Kline & Kline, 2005; Ring, Brener, Knapp & Mailloux, 2015). Meanwhile, many studies utilising the heartbeat discrimination task have documented floor effects as it is the more difficult of the two tasks (Brenner & Ring, 2016). These confounds can generally be mitigated by combining more than one such task, or using appropriate control conditions (Ainley, Brass & Tsakiris, 2014; Tsakiris & Critchley, 2016). Therefore, this study utilised both methods, in addition to a self-report measure of interoceptive accuracy or awareness/sensibility: the MAIA questionnaire (Mehling et al., 2012). We also used a time estimation task as a control for the heartbeat tracking task (Shah, Hall, Catmur & Bird, 2016).

Research utilizing behavioral and self-report methods is necessary to ascertain whether the structural and functional neurological differences found in nicotine-dependent participants translates into observable behavioral differences. Therefore, the aim of the present study was to examine whether a nicotine-addicted population exhibits differences in interoceptive processing using behavioral and self-report methods. Considering the previous findings on interoception and addiction, our first hypothesis was that there would be a difference on measures of interoceptive processing between smokers and non-smokers. Specifically, it was expected that the participants who smoke would demonstrate lower levels of IA and IS. Our second hypothesis was that there would be a relationship between addiction severity and interoceptive processing. Given current neuroimaging evidence (Bi *et* al., 2016; Morales *et al.,* 2014; Stewart *et al.,* 2015) it was expected that there would be a negative relationship between the two constructs: when addiction severity increases, IA and IS decrease.

## Method

### 1. Design

The first level of the experiment was quasi: participants were divided into two groups according to whether they had an addiction to nicotine or not. “To be assigned to the ‘smoker’ condition, participants had to smoke cigarettes daily, and to have been doing so for at least one year. To be assigned to the control condition, participants had to have never had an addiction to smoking, or any other kind of smoking habit. These classifications were determined on the basis of verbal self-reports from participants, and confirmed by scores on the Fagerstrom test. Participants with present or past addictions to any substances other than nicotine were excluded. Participants with present or past addictions to any substances other than nicotine were excluded. Both groups completed the same four measures of IS and IA: two MAIA sub-scales, and two heartbeat perception tasks (see section 3).

The second level of the experiment was correlational. Participants within the ‘smoker’ condition completed the FTND-R (Korte, Capron, Zvolensky & Schmidt, 2013) as a measure of addiction severity, and the relationship between this score and the IS scores from the two MAIA sub-scales and IA scores from the two heartbeat perception tasks was then considered.

### 2. Participants

The sample (N = 100) consisted of 49 cigarette smokers (21 males and 28 females) and 51 non-smokers (19 males and 32 females), aged between 19 and 68 (M = 25.67, SD = 8.71). There were no significant differences in mean age, (p = .730), or resting heartrate (p = .537) between the two groups. The groups were proportionately similar in terms of gender distribution (control condition = 37.3% male, smoker condition = 42.8% male). All participants in the smoker condition abstained from smoking for one hour prior to participation. Overall, the procedure took an average of 30 minutes, therefore it was not deemed necessary to take measures of nicotine withdrawal.

All participants gave written informed consent. Participants were offered payment at a rate of £7 per hour or participation credits. The protocol was approved by the Psychology Departmental Research Ethics Panel (DREP) and ratified by the Faculty Research Ethics Panel in November (2015) under the terms of Anglia Ruskin University’s Policy and Code of Practice for the Conduct of Research with Human Participants.

### 3. Apparatus and Procedure

#### 3.1 Multidimensional Assessment of Interoceptive Awareness questionnaire (MAIA; Mehling *et al*., 2012)

The MAIA is a self-report measure of IA that comprises eight inter-related sub-scales that provide a multi-dimensional profile of interoceptive processing, with each sub-scale assessing a different facet of IA. All participants completed the two sub-scales which were deemed most relevant to the hypothesis: the ‘noticing’ and ‘attention regulation’ scales. The ‘noticing’ scale consists of four items, and was selected because it measures the subjective perception of the ability to perceive and focus on bodily sensations, specifically testing awareness of comfortable, uncomfortable and neutral sensations. Questions include, ‘When I am tense I notice where the tension is located in my body’. Meanwhile, the ‘attention regulation’ scale consists of seven items and measures the ability to control and maintain attention towards bodily sensations. Questions include, ‘I can pay attention to my breath without being distracted by things happening around me.’ Answers for both scales are given on a 6-point Likert scale, ranging from ‘never’ to ‘always’. In the present study, Cronbach’s α was 0.70 for the ‘noticing’ scale, and 0.82 for the ‘attending’ scale, which is comparable to previous assessments (Mehling et al., 2012), and indicates good levels of internal consistency reliability (Kline, 2005).

### 3.2 Revised Fagerström Test for Nicotine Dependence (Korte *et al.*, 2013)

The FTND-R was completed by participants within the ‘addiction’ condition as a measure of the severity of addiction to smoking. The FTND-R has moderate levels of validity and reliability, and significantly predicts bio-chemical measures of cigarette-smoking addiction, such as carbon monoxide (CO) levels in the bloodstream (Korte *et al.,* 2013). It is comprised of six questions with responses recorded on 4-point Likert scales. Questions include, ‘How many cigarettes a day do you smoke?’ In the present study, Cronbach’s α was 0.69, which is homogenous with previous findings (Korte *et al.,* 2013) and indicates appropriate internal consistency reliability (Kline, 2005).

#### 3.3 Heartbeat tracking task (Schandry, 1981)

Chart 5 software (Windows) and a Powerlab data acquisition unit (ADInstruments, Germany) were connected to a PC to measure cardiac events during the heartbeat tracking task. Three disposable electrocardiography (ECG) electrodes and conductive hydrogel (positioned on the chest in standard three-lead configuration) relayed R-wave output through shielded wires. The electrodes were self-attached by participants under their clothes, using a visual diagram. Throughout this task, participants were seated in an upright position and asked to attempt to sense their heart beating from the inside of their body, without using a manual pulse. Once they had identified a sensation that they felt indicated their heart beating, participants were asked to count the number of heartbeats occurring in four discrete time intervals of 25, 35, 45, and 55 seconds (presented in a random order across participants). The beginning and end of each trail was signified by an auditory start/stop cue. Participants were unaware of how long they were counting for, and no performance-related feedback was given.

#### 3.4 Time estimation task (Shah *et al.,* 2016)

Performance on the heartbeat tracking task might be confounded by one’s ability to judge the duration of time intervals, and some evidence suggests that smokers may have poorer time estimation abilities than non-smokers, particularly during periods of abstinence (Ashare & Kable, 2015; Klein, Corwin & Stine, 2002; Sayette, Loewenstein, Kirchner & Travis, 2005). We controlled for this by asking half of the participants (randomly selected) in both conditions (*n* = 50) to complete a time estimation task. Participants were asked to estimate elapsed time during three discrete time intervals of 19, 37, 49 seconds (presented in a random order across participants). Again, participants were unaware of how long they were counting for, and no performance-related feedback was given.

#### 3.5 Heartbeat discrimination task (Brener & Kluvitse, 1988; Katkin *et al.*, 1983; Whitehead *et al.*, 1977)

This task was executed using software developed in-house (http://lnco.epfl.ch/expyvr) and the same disposable ECG electrodes, which were connected to a laptop with built-in speakers. Participants had to listen to a series of tones **(lasting 100 ms each)** and state verbally to the experimenter whether they thought the tones in each trial were synchronous or asynchronous to their own heartbeat. For the synchronous condition, the software produced a tone that was synchronous with the R-wave of the QRS complex of the ECG to create a rhythm that was concurrent to the participant’s own heartbeat. The software computed, at 60Hz, the instantaneous derivative of the ECG signal (buffered data) to detect the high-amplitude signal change between the Q and R peaks of the ECG. Continuous adjustment of the algorithm to cumulatively averaged extrema allowed for an automatic adaptation to inter-participant differences and signal amplitude variations. The imprecision of the R-peak detection in the software is maximally 1 frame (33ms). As the beep has a fixed duration of 100 ms, the ‘mid-point’ of the beep therefore occurs on average at 73 +/-16 ms after the R peak. This is therefore close to synchronous with the theoretical blood flow at the aortic root, and thus with the systolic heart contraction. Previous research has shown that participants do not feel their heartbeats as synchronous with an external stimulus unless there is a delay of around 100-250 ms between it and the R-peak of the ECG (Brener & Kluvitse, 1988; Wiens and Palmer, 2001). For the asynchronous condition, the software produced a tone that was either 80% or 120% of the speed of the two preceding R-waves to create a rhythm that was asynchronous to the participant’s own heartbeat. In this condition the system computes, online, a running average of the participant’s ECG frequency. This frequency is then continuously used as a reference for computing the timing of the beeps (80% or 120% of the ECG frequency). The timing between the peaks of these two out-of-phase signals of different frequency (e.g. for a participant’s ECG at 72 beats per minute, the beeping at 80% will be at a rate of 57.6 BPM) is therefore continuously changing between zero and half of the period of the faster of the two signals (i.e. between 0 and 347 ms in the above example). Generating asynchrony in this dynamic way therefore precludes the possibility of the beeps and R-peaks being synchronous and is therefore better than simply employing a fixed delay between the two signals (which would correspond to a constant phase shift and could, for participants with HRs of certain frequencies, lead to the asynchronous condition being synchronous). Each trial was comprised of 20 tones. Participants were presented with 16 trials, of which eight were synchronous and eight were asynchronous (ordered randomly). The eight asynchronous trials were divided into four that were at 80% of the participant’s R-wave signal rate, and four that were at 120%, also with a random order. Each participant had one practice trial to allow for familiarization with the procedure (the synchronicity of this trial was assigned randomly). Participants completed the task without the use of a manual pulse, and no performance-related feedback was given. An identical task was used in recently published papers by Piech *et al.* (2017) **and Mul et al., (2018)**

## Results

### 4.1 Transformation of raw data

Three separate processes were used to transform the raw data from each task into IS and IA scores that could be analyzed. Firstly, an IS score for the heartbeat tracking task was calculated by comparing the number of cardiac events that participants counted with the number of R-waves observed for each of the four trials, in accordance with the formula: 1⁄4 ∑ (1 – [recorded heartbeats – counted heartbeats]/recorded heartbeats), as used by Schandry (1981). Absolute values are utilised, so that scores range from 0 to 1, those closer to 1 indicate higher IS. Similarly, the time estimation task scores were computed as percentage accuracies, in accordance with the formula: 1/3 ∑ (1 – [actual elapsed time – estimated elapsed time]/actual elapsed time), as used by Shah *et al.* (2016). Secondly, an IS score for the heartbeat discrimination task was calculated by dividing the number of correct answers by the total number of trials to generate a percentage accuracy score. Scores again range from 0 to 1, with those closer to 1 indicating higher IS. Finally, two IA scores were recorded for each participant, with ‘noticing’ and ‘attention regulation’ abilities represented by the mean response from each scale respectively. Scores range from 0 to 5, those closer to 5 indicate higher IA.

### 4.2 Testing hypothesis one

Data from the heartbeat tracking and discrimination tasks and the MAIA ‘attending’ scale met assumptions of normality. Therefore, independent-samples *t*-tests were used to determine if there were differences in IS and IA scores between the ‘smoker’ and ‘non-smoker’ conditions. Because of the multiple comparisons (*k*), a Bonferroni adjustment was applied to reduce the chance of Type I error, such that *p* = (1 – α)^*k*^ ≈ 1 – *k*α = α/*k* = .01. For the heartbeat tracking task, non-smokers (0.74 ± 0.15) performed better than smokers (0.65 ± 0.18), a statistically significant difference of 0.09 (95% CI, 0.03 to 0.15), *t*(98) = 2.77, *p* = 0.007, *d* = 0.55 (see Figure 1). For the heartbeat discrimination task, non-smokers (0.60 ± 0.16) performed better than smokers (0.56 ± 0.17), but the difference of 0.04 (95% CI, -0.02 to 0.11) was not statistically significant, *t*(98) = 1.26, *p* = 0.209, *d* = 0.25. Similarly, while non-smokers (3.01 ± 0.77) produced higher IA ratings than smokers (2.94 ± 0.74), the difference in scores from the MAIA ‘attending’ scale (0.67; 95% CI, -0.23 to 0.37) was not significantly different between groups *t*(98) = 0.445, *p* = 0.657, *d* = 0.09.

**Figure 1.**
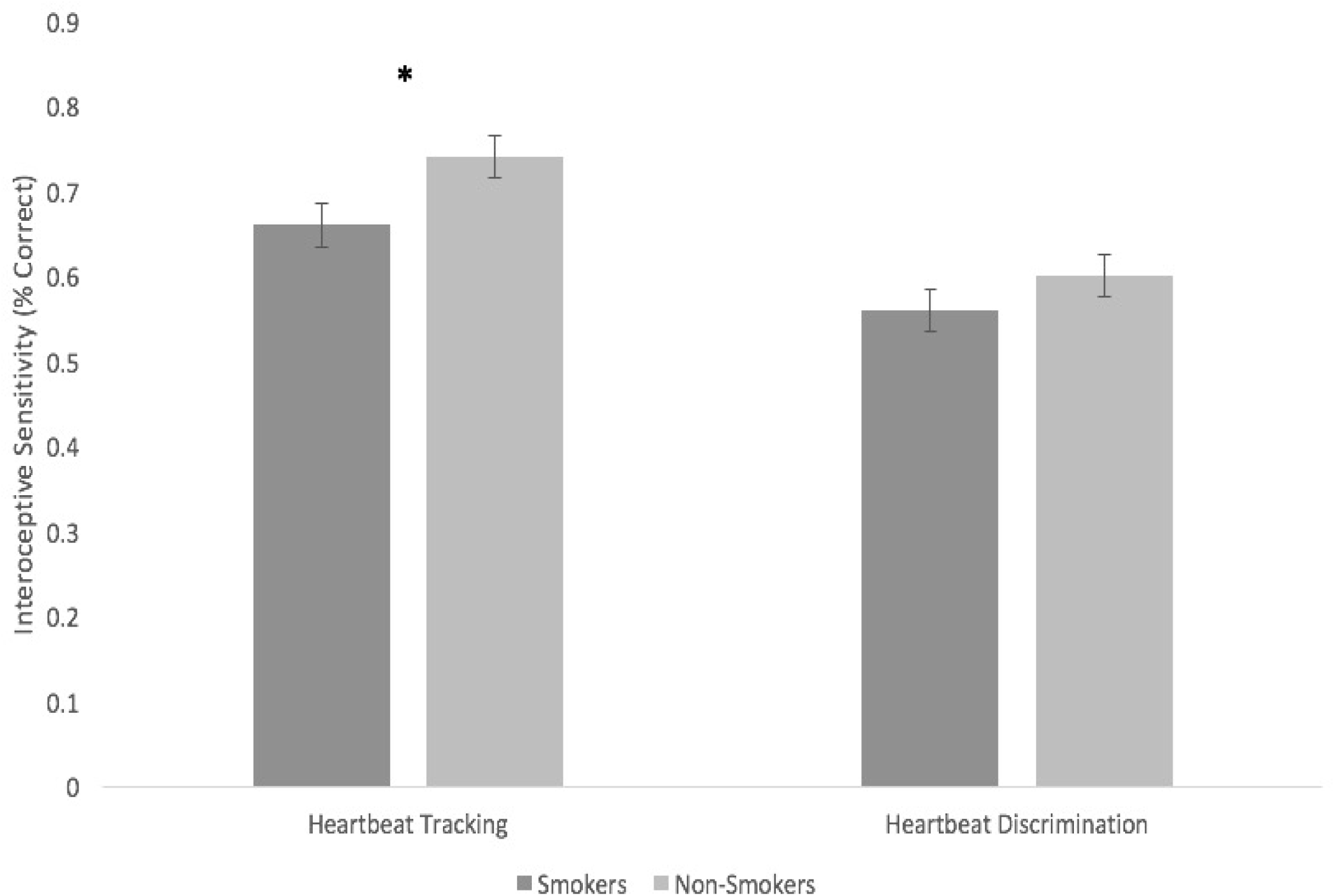
Bar chart with standard error bars to show group differences in interoceptive sensitivity, as assessed by the heartbeat tracking and heartbeat discrimination tasks.

There were outliers in scores from the MAIA ‘noticing’ scale, and the distribution for the ‘no addiction’ condition was skewed and Kurtotic(the z-scores for skewness and kurtosis were 4.14 and 3.36, respectively). Therefore, a Mann-Whitney U test was run to determine if there were differences in IA scores from the MAIA ‘noticing’ scales between the ‘addiction’ and ‘no addiction’ conditions. Median ‘noticing’ scores were not statistically different between smokers (*Mdn* = 3.50) and non-smokers (*Mdn* = 3.50), *U* = 1228.0, z = -.149, *p* = .881.

There were also outliers in the time estimation data, and the distribution was skewed and kurtosed for both conditions (for the ‘no addiction’ condition, the z-scores for skewness and kurtosis were 3.38 and 3.08, respectively, and for the ‘addiction’ condition the z-scores for skewness and kurtosis were 3.45 and 3.15). Therefore, a Mann-Whitney U test was run to determine if there were differences in time estimation accuracy scores between the ‘addiction’ and ‘no addiction’ conditions. Whilst the ‘no addiction’ condition (*Mdn* = 82.59) demonstrated greater accuracy in comparison to the ‘addiction’ condition (*Mdn* = 77.15), the difference between the groups was not statistically significant after Bonferroni correction *U* = 187.0, z = -2.435, *p* = .015. Furthermore, time estimation was not significantly correlated with IS scores from the heartbeat tracking task for the both the ‘addiction’ and ‘no addiction’ conditions; *r*_*s*_ (23) = .215, *p* = .302; *r*_*s*_ (23) = .162, *p* = .440, respectively. This suggests that the difference between smokers and non-smokers on the heartbeat tracking task was not due to differences in time estimation accuracy.

### 4.3 Testing hypothesis two

To assess whether addiction severity was related to IS and IA, we computed Spearman’s rank-order correlations between the FTND-R and the IA and IS scores from the heartbeat perception tasks and the MAIA scales for the ‘addiction’ condition (*n* = 49). As can be seen in Table 1, FTND-R scores were not correlated with IS scores from the heartbeat tracking task, *r*_*s*_ (47) = .179, *p* = .218, or the heartbeat discrimination task, *r*_*s*_(47) = .188, *p* = .197. Similarly, FNTD-R scores were not correlated with IA scores from the MAIA ‘noticing’ *r*_*s*_(47) = .160, *p* = .271; or ‘attending’ scales, *r*_*s*_(47) = .188, *p* = .197 (see Figure 2).

**Table 1.** Correlations between all variables, with results for the ‘Addiction’ in the top diagonal and for the ‘No addiction’ condition in the bottom diagonal

|  | (1) | (2) | (3) | (4) | (5) | (6)<sup>+</sup> | (7) |
| --- | --- | --- | --- | --- | --- | --- | --- |
| (1) Heartbeat Tracking |  | .015 | .183 | -.004 | .179 | .215 | .531** |
| (2) Heartbeat Discrimination | .278* |  | .087 | -.023 | .188 | .548** | .326* |
| (3) MAIA ‘Noticing’ | -.082 | .165 |  | .421** | .160 | -.032 | -.224 |
| (4) MAIA ‘Attending’ | -.137 | .053 | .523** |  | .188 | -.180 | -.188 |
| (5) FTND-R | -- | -- | -- | -- |  | -.155 | -.151 |
| (6) Time Estimation<sup>+</sup> | -.162 | .378 | -.036 | .099 | -- |  | .064 |
| (7) Average Heartrate | -.213 | -.127 | -.006 | .092 | -- | -.105 |  |

*Note.* ‘Addiction’ condition $n = 49$ , ‘No addiction’ condition $n = 51$ . \* $p < .05$ , \*\* $p < .001$ . <sup>+</sup> $n = 25$ for both conditions

**Figure 2.**
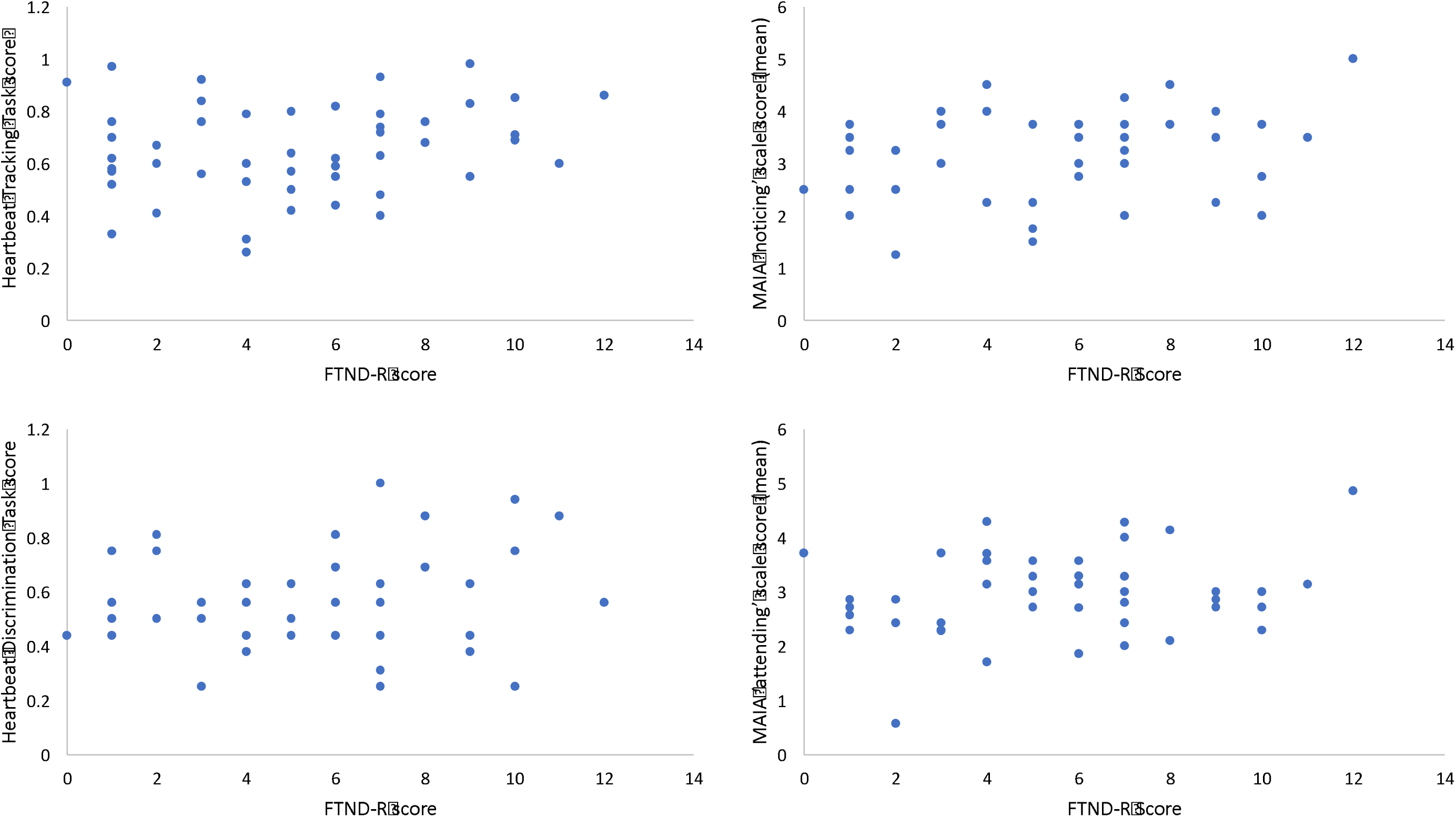
Scatter plots to show correlations between the FTND-R and the heartbeat perception tasks and MAIA scales (n = 49).

## Discussion

The current study sought to examine the associations between smoking and interoceptive sensitivity (IS) and interoceptive awareness (IA). We found that the IS of participants who had never smoked was greater than that of participants with an addiction to smoking cigarettes, but there were no significant group differences in IA. Specifically, participants in the control group performed significantly better on the heartbeat tracking task than participants in the ‘smoker’ group. However, no significant differences between groups were observed for the heartbeat discrimination task, or the MAIA measures of IA. Meanwhile, within the 'smoker' group, neither IS nor IA were associated with addiction severity.

The observed difference in IS between the ‘smoker’ and ‘non-smoker’ groups is the first behavioural demonstration of such a difference in a population addicted to smoking. It is consistent with our hypothesis that addiction to smoking cigarettes would be associated with lower IS. The finding also aligns with recent studies, which point towards a link between addiction and deficits in IS (Ates Col *et al*., 2016; Sönmez *et al.,* 2016), and supports previous research demonstrating that in addiction populations, some interoceptive brain areas show functional differences (Avery *et* al., 2016; Bi *et* al., 2016; Droutman *et al.,* 2015) and have structural abnormalities (Goldstein *et* al., 2009; Morales *et al.,* 2014). With support growing for the hypothesis that interoceptive processing may be disturbed in addiction (Ates Col *et al*., 2016; Sönmez *et al.,* 2016), it has been argued that an enhanced conceptualisation of addiction should include compromised interoception, alongside other addiction-relevant constructs (Goldstein *et* al., 2009, Verdejo-Garcia, Clark & Dunn, 2012).

However, a crucial question remains: how does compromised interoceptive processing relate to addiction? Verdejo-García and Bechara (2009) integrate the evidence for disturbed interoception and altered decision-making processes in substance addicts, to explain the underlying mechanisms of substance addiction using the Somatic Marker Hypothesis (Damasio, 1996). According to this theory, complex decision making processes are contingent upon emotion-based bodily signals **–** ‘somatic markers’ **–** which are processed in structures such as the ventromedial prefrontal cortex, the insula, and the amygdala. It may be that structural and functional differences in regions such as the insula render substance addicts less sensitive to bodily signals conveying the negative consequences of certain courses of action; which in turn increases the probability of engaging in risky drug-taking behaviours (Verdejo-García & Bechara, 2009; Verdejo-Garcia *et al*., 2012). The apparent identification of deficits in IS for smokers in the current study corroborates this theory.

Whilst we identified differences between smokers and non-smokers on the heartbeat tracking task, no such differences were identified for the self-report (MAIA) measures. This is not a unique finding; a poor correspondence between behavioural (heartbeat perception tasks) and self-report measures of interoceptive processing has been previously identified (Garfinkel *et al.,* 2015; Forkmann *et al*., 2016). Indeed, Garfinkel *et al.* (2015) present empirical support for a dissociation between IS and IA, which they conceptualised as two of three distinct constructs within interoceptive processing: Interoceptive *accuracy* (objective measures of the perception of internal bodily states; also termed ‘interoceptive sensitivity’); interoceptive *sensibility* (subjective, self-reported perceptions; also referred to as ‘interoceptive awareness’); and a metacognitive measure of overall interoceptive processing (quantifying the relationship between subjective interoceptive awareness and objective interoceptive sensitivity; which was not measured in the current study). This three-dimensional model of interoception is underpinned by evidence of distinct patterns of functional connectivity in relation to objective performance and subjective beliefs (Barttfield *et al.,* 2013). In reference to the present study, it could be that group differences in performance on the objective heartbeat tracking task reflect underlying group differences in areas associated with heartbeat perception, such as the right anterior insula (Critchley *et al.* 2004), whilst areas underlying the subjective interpretation of such bodily signals, (for example, the orbitofrontal areas; Flemming, 2012) could remain unaffected, explaining the lack of group differences in IA.

The present study utilized two heartbeat perception tasks, and whilst significant group differences were identified using the heartbeat tracking task, no such differences were identified using the heartbeat discrimination task. We did find a correlation between the two tasks, but only for non-smokers (see Table 1), suggesting that the tasks assess a similar facet of IS for non-smokers, but not for smokers. Similarly, scores for the heartbeat tracking task were significantly positively correlated with resting heart rate for smokers, whilst they were negatively (but not significantly) correlated with task scores for non-smokers (Table 1). The positive correlation between heart rate and task performance in smokers contrasts with current research, which suggests that a negative correlation between the two variables is typical (e.g. Zamariola, Maurage, Luminet & Corneille, 2018). Whilst resting heart rates of the smokers (*M* = 73.6 beats per minute) tended to be higher than the non-smokers (*M* = 71.9 beats per minute), there were no statistically significant differences between the two groups (*p* = .537). Given the lack of statistically significant difference, and that the performance of participants in the smoker condition tended to be worse than that of participants in the control condition, the positive correlation between resting heart rate and task performance for smokers is surprising. One possible explanation for the results is that participants varied in their knowledge of their average resting heart rates, a factor that has been shown to confound tracking task performance (Ring, Brenner, Knapp & Mailloux, 2015). It is possible that non-smokers may have a greater knowledge of their resting heart rate, which could enable them to estimate their heart rate more accurately during the tracking task, and that the smokers with higher heart rates also tended to have a greater knowledge of their average resting heart rates. These are avenues that should be pursued in future research.

Whilst some previous studies have found correlations between the heartbeat perception tasks (Hart, McGowan, Minati & Critchley, 2013; Knoll & Hodapp, 1992) others have not (Betka *et al.,* 2018; Forkmann *et al*., 2016; Michal *et al*., 2014; Schulz, Lass-Hennemann, Sütterlin, Schächinger & Vögele, 2013). The tasks are assumed to measure IS in fairly equivalent ways, but actually require quite distinct psychological processing (Forkmann *et al*., 2016). The heartbeat tracking task requires some sense of the duration of time intervals, which we controlled for with the time estimation task. We found no difference in performance on the time estimation task between smokers and non-smokers, however this was only tested in a sub-set (half) of the sample this was only tested in a sub-set (half) of the sample and the difference was close to being significant, and we acknowledge the limitation of this. , The heartbeat discrimination task, on the other hand, requires a comparison of the timing of events across two different sensory modalities (cardiac interoceptive and auditory). For this reason, the heartbeat discrimination task is generally considered to be a harder and very different task, with most participants performing close to chance level (Brenner & Ring, 2016; Knapp-Kline & Kline, 2005; Phillips, Jones & Snell, 1999). Indeed, several other studies have identified group differences for the heartbeat tracking task, but not for the heartbeat discrimination task (e.g. Kandasamy *et al.,* 2016; Mul, Stagg, Herbelin & Aspell, 2018).

Our second main finding did not support our prediction: IS and IA were not found to be significant predictors of addiction severity. This aligns with some recent research (Sönmez *et al*., 2016), but contrasts with other research where a negative relationship between interoception and addiction was identified (Bi *et al.,* 2016; Stewart *et al*., 2015). It is important to note that the dataset was divided in half for the correlational analyses. Therefore, the result may be a reflection of low statistical power. Moreover, there was little variance in data for the FTND-R, which limits the potential for identifying associations between addiction severity and IS and IA: scores on the FTND-R range from 0 to 16, but in our dataset there were no scores greater than 12. Future studies could attempt to recruit participants with greater levels of addiction severity, or use alternative assessments such as CO levels which may produce more nuanced data.

Clinically, the observation of decreased IS in smokers could lead to new treatment approaches aiming to alter awareness of interoceptive signals. This is important because current national smoking cessation programmes are associated with modest results (Centre HSCI, 2014). For example, cessation programs could be enhanced by including therapies aimed at improving awareness of the body, such as biological feedback education, body-focused meditations, and mindfulness interventions (Farb, Segal & Anderson, 2013; Paulus, Stewart & Haase, 2013; Verdejo-Garcia *et al.,* 2012). Indeed, pilot research suggests that the use of a Mindful Awareness in Body-Oriented Therapy intervention was viewed by addicts as an essential component of relapse prevention (Price & Smith-DiJulio, 2016). There is also potential for the use of biological measures of IS, such as the heartbeat evoked potential (HEP; Pollatos, Kirsch & Schandry, 2005) being used as a biomarker of substance use disorders.

Future research is also required to clarify whether additional variables that could make one more likely to smoke cigarettes and perform poorly on the heartbeat tracking task (for example, trait anxiety or impulsivity) could be responsible for the observed association between smoking and tracking task performance in the present study. Furthermore, work is also required to ascertain the causal direction of the relationship between interoception and addiction, which was unidentifiable both in the present study, and within previous studies on the topic, given the correlational nature of the research designs utilised. A cue-induced craving paradigm, longitudinal design, or the addition of a group of participants who no longer meet the criteria for addiction to cigarette smoking may facilitate clarification.

## Acknowledgements

Farah Hina is grateful to the Wellcome Trust for providing support in the form of a Biomedical Vacation Scholarship (grant number: 206882/Z/17/Z)

